# Multiscale Entropy Analysis of Retinal Signals Reveals Reduced Complexity in a Mouse Model of Alzheimer’s Disease

**DOI:** 10.1101/2022.01.27.478055

**Authors:** Joaquín Araya-Arriagada, Sebastián Garay, Cristóbal Rojas, Claudia Duran-Aniotz, Adrián G. Palacios, Max Chacón, Leonel E. Medina

## Abstract

Alzheimer’s disease (AD) is one of the most significant health challenges of our time, affecting a growing number of the elderly population. In recent years, the retina has received increased attention as a candidate for AD biomarkers since it appears to manifest the pathological signatures of the disease. Therefore, its electrical activity may hint at AD-related physiological changes. However, it is unclear how AD affects retinal electrophysiology and what tools are more appropriate to detect these possible changes. In this study, we used entropy tools to estimate the complexity of the dynamics of healthy and diseased retinas at different ages. We recorded microelectroretinogram responses to visual stimuli of different nature from retinas of young and adult, wild-type and 5xFAD –an animal model of AD– mice. To estimate the complexity of signals, we used the multiscale entropy approach, which calculates the entropy at several time scales using a coarse graining procedure. We found that young retinas had more complex responses to different visual stimuli. Further, the responses of young, wild-type retinas to natural-like stimuli exhibited significantly higher complexity than young, 5xFAD retinas. Our findings support a theory of complexity-loss with aging and disease and can have significant implications for early AD diagnosis.

## Introduction

Alzheimer’s disease (AD) is a chronic neurodegenerative disorder and the most common form of dementia, affecting between 10 to 30% of the population >65 years old^1^. The symptoms of AD progress from mild memory loss to severe dementia, include language difficulties, psychiatric symptoms, behavioral and visual disturbances, and difficulties performing memory and motor daily living activities^2^. Although the causes of AD are unknown and its pathophysiological mechanisms only partially understood, two pathological key signatures of the disease have been identified: the accumulation of a nonsoluble form of amyloid-*β* peptides in extracellular spaces and in blood vessels, and the presence of neurofibrillary tangles in neurons due to the aggregation of protein tau^3^. Early diagnosis of AD can be a challenging task because degeneration of neurons in the cortex and/or the hippocampus begins years before the manifestation of the symptoms, and a definitive diagnosis can only be established post-mortem^4^. Amyloid-*β* and/or protein tau accumulation can be detected utilizing positron emission tomography (PET) imaging or cerebrospinal fluid (CSF) analysis^1^, but these techniques are invasive and often costly. In recent years, the ocular globe has received increased attention as a candidate expressing AD biomarkers due to the mounting evidence of eye abnormalities observed in AD patients, including retinal thinning, optic nerve damage, degeneration of retinal ganglion cells, and changes to vascular parameters^4–6^. Further, the retina of AD patients appears to accumulate amyloid-*β* and protein tau, and this manifestation may be related to the visual difficulties that these patients exhibit^5, 7–10^.

Functional changes in the retina of AD patients can be detected using the electroretinogram (ERG)^11–15^, in which electrodes placed on the cornea or on the skin beneath the eye are used to record the bio-electrical activity of photoreceptors, bipolar cells, and other neurons of the retinal network^16, 17^. Although normal ERG recordings in AD patients have been reported in the past^18^, recent evidence has demonstrated amplitude reductions and latency increments in ERG responses of patients at different stages of the disease^4, 19^. As well, similar findings have been obtained in animal studies of neurodegeneration. For example, ERG measurements in the transgenic 5xFAD mouse, an animal model of AD, exhibited abnormalities that accentuated with age^20^, and *ex vivo* multi-electrode array (MEA) recordings from the 5xFAD’s retinal ganglion cells (RGCs) showed alterations in the firing activity that appeared to arise from a neurochemical imbalance in this animal model^21^. Taken together, these findings suggest that the early visual bio-electrical activity of the retina can hint at AD-related physiological changes, but further evidence needs to accrue, and additional signal analysis tools need to be explored.

In the last two decades, there has been growing interest in entropy and other information theory tools for biological signal analysis after the introduction of the multiscale entropy (MSE) method^22^. Under this framework, a measure of the entropy of a physiological output is calculated for multiple time scales, resulting in a MSE curve. Then, the MSE curve provides an estimation of the degree of complexity of the physiologic system. In this case, entropy refers to signal disorder, and complexity, to the nonlinear interactions between various structural units and their collective behavior, operating over a wide range of temporal and spatial scales^23^. Physiological complexity has received increased attention due to the theory of complexity-loss with aging and disease^23–26^. This theory postulates that, as stated by Goldberger *et al*.^24^: 1) “the output of healthy systems (…) reveals a type of complex variability associated with long-range correlations”, and 2) “this type of multiscale, nonlinear complexity breaks down with aging and disease, reducing the adaptive capabilities of the individual”. Recent evidence seems to support this theory as, for instance, entropy measures of cerebral blood flow velocity decayed in altered states^27^, actigraphy data exhibited lower complexity in depressed subjects as compared to non-depressed controls^28^, and complexity of EEG signals from subjects with AD correlated well with performance in working memory tasks^29^. MSE analysis can have significant implications in biomedical research as several studies have demonstrated its application to detection of heartbeat irregularity^22, 30^, postural control alterations^31^ and postoperative neurocognitive dysfunction^32^, and diagnosis of depression^28^, Parkinson’s disease^33^, schizophrenia^34^, and AD^35^.

In this study, we performed MEA analyses of μERG signals to compare the electrophysiological behavior of healthy (WT) and diseased (5xFAD) retinas of young and adult subjects, in response to synthetic and natural-like visual stimuli. In agreement with the complexity-loss theory, we found that the retinas of younger animals exhibit higher entropy levels across a wide range of MSE, revealing a more complex pattern of physiological activity than that of older retinas, and that for certain inputs, the young retina of a mouse model of AD expresses a less complex network activity than that of a healthy one. These findings can have great implications in the search for biomarkers for early AD diagnosis.

## Results

### Time- and frequency-domain characteristics of microelectroretinogram recordings

We recorded the microelectroretinogram (*μ*ERG) from pieces of retina of wild-type (WT) and 5xFAD mice in response to visual inputs during two stimulation protocols, namely, the chirp stimulus (CS) protocol and the natural image (NI) protocol (Fig. 1). The retinal responses showed great consistency across trials and subjects. For the CS protocol, clear onset and offset responses were observed for the steps of light applied in the initial ~ 6 s (Fig. 2a). Further, the *μ*ERG showed an oscillatory response that followed the increase in the frequency of the sinusoidal light stimulus, albeit with decreasing signal amplitude, in agreement with previous reports that showed reduced *μ*ERG amplitude for stimulus frequency beyond ~ 10 Hz^11^. The responses recorded for the NI protocol exhibited higher variability across subjects (Fig. 3a), and a clear onset response in the transition from dark to light that resembled that for the CS protocol. For both CS and NI protocols, the power spectral density (PSD) was similar across the four groups (Figs. 2b and 3b), albeit the 5xFAD-young group had somewhat higher power in the 10 – 15 Hz band. In addition, the PSD showed a peak at a very low frequency (~ 2 Hz). The coherence between the CS input signal and the *μ*ERG response was nonzero for frequencies ≤ 15 Hz, with a peak at ~ 8 Hz, and was somewhat higher for young retinas in the 8-15 Hz range (Fig. 2c). In addition, *μ*ERG signals did not show any salient feature among the groups for amplitude-based variability metrics like the mean and the standard deviation (Fig. 3c). In summary, the time- and frequency-domain characteristics of the *μ*ERG responses did not show any notable difference due to condition (WT vs 5xFAD) or age (young vs adult).

**Figure 1.**
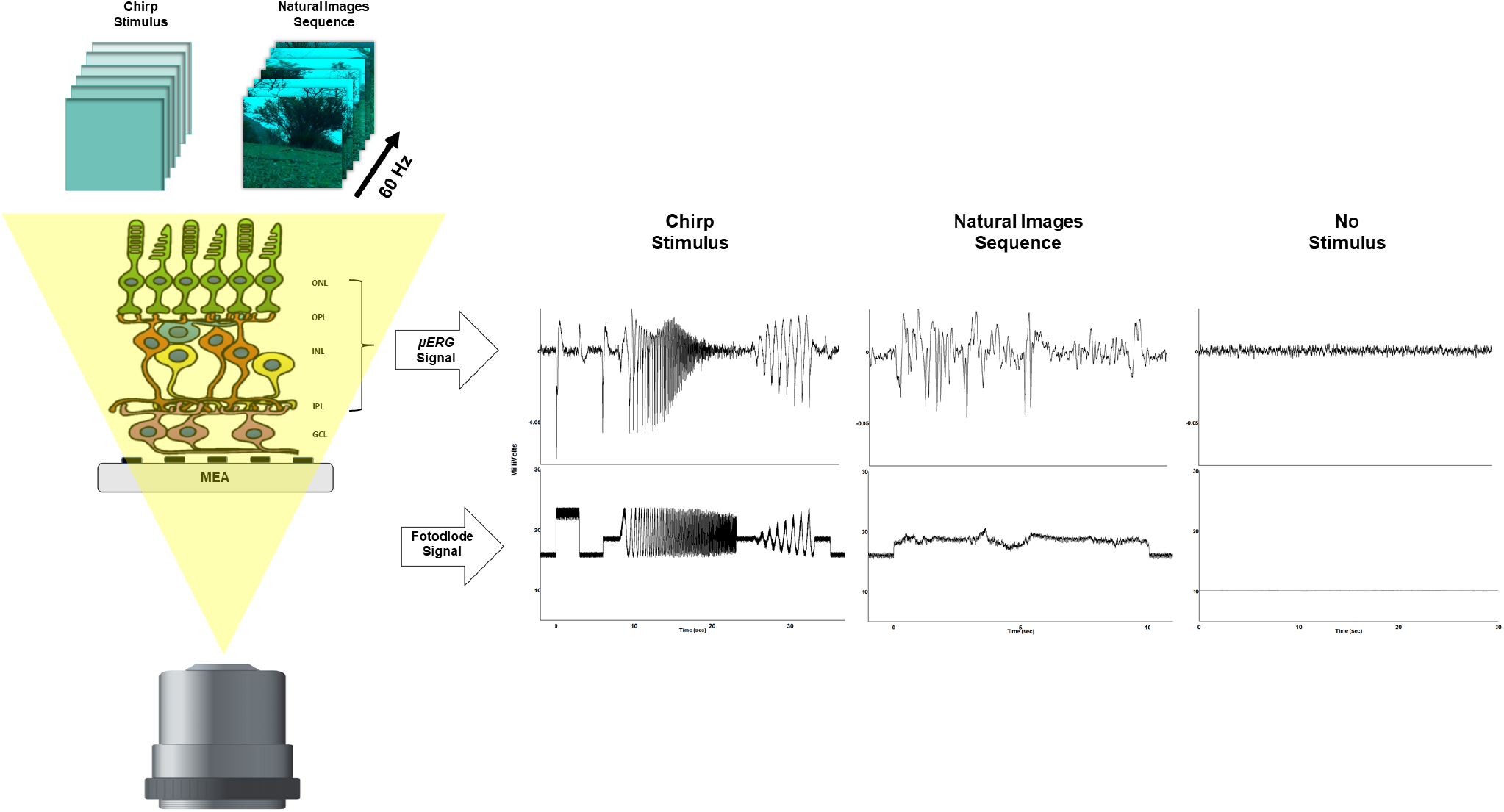
Multielectrode array (MEA) recordings of retinal responses to visual stimuli. A MEA was mounted on a piece of retina to record *μ*ERG signals of the electrophysiological activity of several cell types of the retinal network, including those in the outer nuclear layer (ONL), outer plexiform layer (OPL), inner nuclear layer (INL) and inner plexiform layer (IPL). The firing activity of retinal ganglion cells (GCL) was low-pass filtered of the *μ*ERG. Two types of visual stimuli were applied using a LED projector: 1) A so-called chirp stimulus, and 2) A sequence of natural images. Examples of peristimulis time histogram (PSTH) from the *μ*ERG recordings, in response to 21 repetitions of the chirp and the natural image (on the right side), together with a fotodiode signal recorded simultaneously to keep track of the stimulus intensity. For reference, it is also shown a *μ*ERG recording during a period of no visual stimulation.

**Figure 2.**
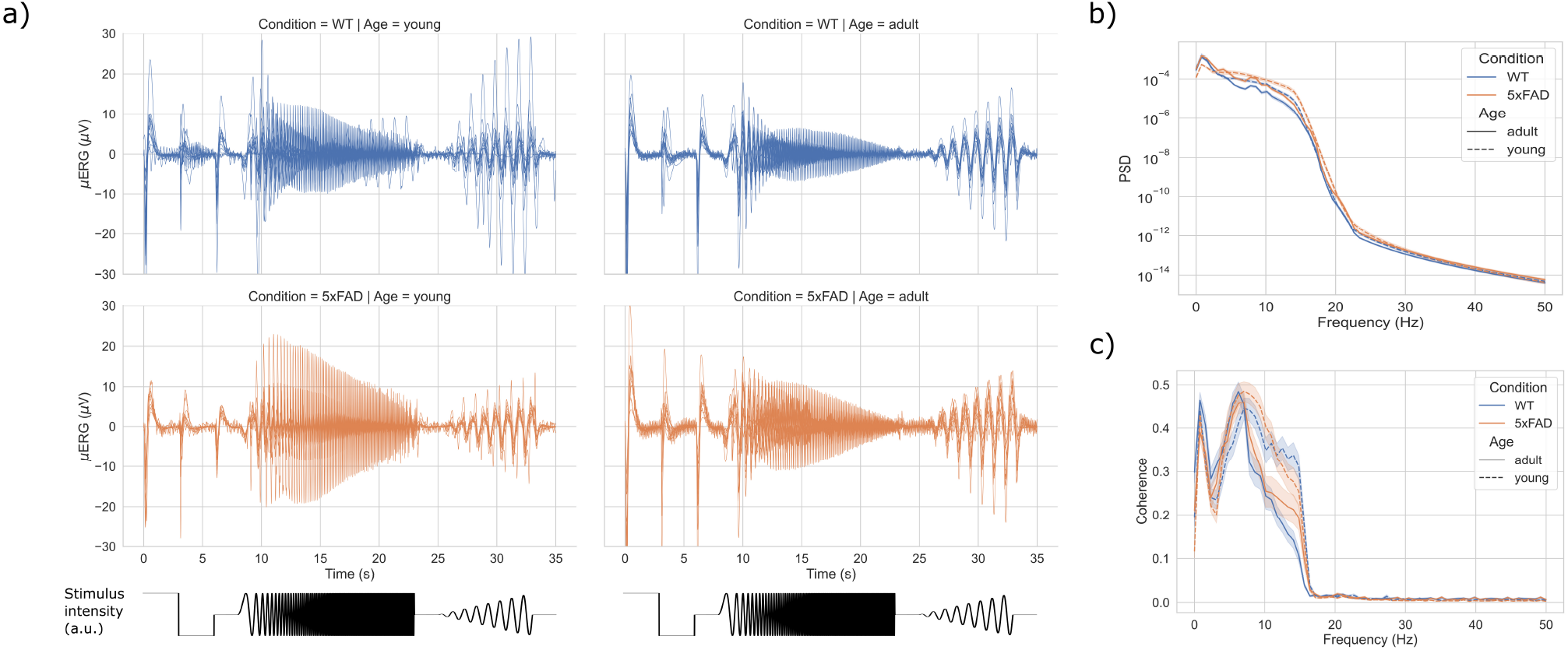
Time- and frequency-domain characteristics of *μ*ERG responses during the CS protocol. a) *μ*ERG responses during the CS protocol for all retinas separated by condition and age. For reference, the stimulus intensity is shown on the bottom. b) Average power spectral density (PSD) of the *μ*ERG responses. c) Coherence between the light stimulus and the *μ*ERG response for the CS protocol. Shaded areas indicate 95% confidence intervals.

**Figure 3.**
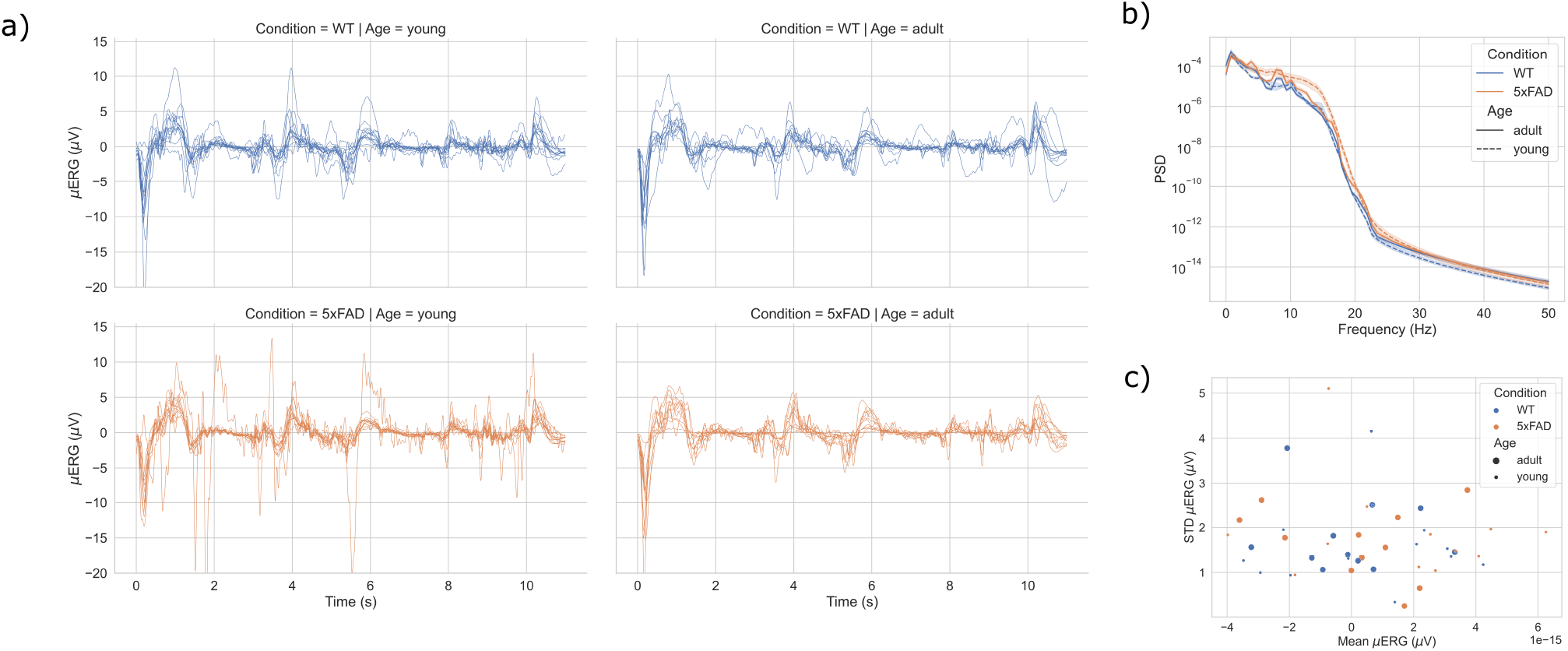
Time- and frequency-domain characteristics of *μ*ERG responses during the NI protocol. a) *μ*ERG responses during the NI protocol for all retinas separated by condition and age. b) Average power spectral density (PSD) of the *μ*ERG responses. c) Average signal and standard deviation of *μ*ERG responses. Shaded areas indicate 95% confidence intervals.

### Multiscale entropy analysis

For the multiscale entropy (MSE) analyses, we first calculated the fuzzy entropy (*FuzzyEn*)36 of the *μ*ERG signals at different time scales, thereby obtaining a MSE curve. Next, we estimated a complexity index for each retina as the area under the MSE curve^37^. The entropy of the *μ*ERG signals increased rapidly for small scales of up to 10 (Figs. 4a and 5a). Above this scale, the entropy for the CS protocol stabilized around ~ 0.6 for young retinas and ~ 0.5 for older retinas (Figs. 4a). This resulted in higher complexity – as quantified as cumulative entropy for scales from 20 to 44 – for young retinas of WT and 5xFAD subjects (Figs. 4b), and the median (± 25-75% interquartile range) complexity index for WT-young, 5xFAD-young, WT-adult and 5xFAD-adult was 14.7 (± 3.85), 15.04 (± 3.82), 12.82 (± 3.25) and 11.33 (± 2.43), respectively (Table 1). For the NI protocol, the entropy increased monotonically as the scale increased for all retinas (Fig. 5a). Further, the average MSE was consistently higher for WT retinas as compared to 5xFAD ones, and as well, it was consistently higher for young retinas as compared to adult ones. Consequently, the complexity of retinal responses was higher for WT-young retinas, and the median (± 25-75% interquartile range) complexity index for WT-young, 5xFAD-young, WT-adult and 5xFAD-adult was 22.72 (± 3.21), 21.26 (± 2.14), 21.22 (± 5.32) and 19.92 (± 5.63), respectively (Table 1).

**Figure 4.**
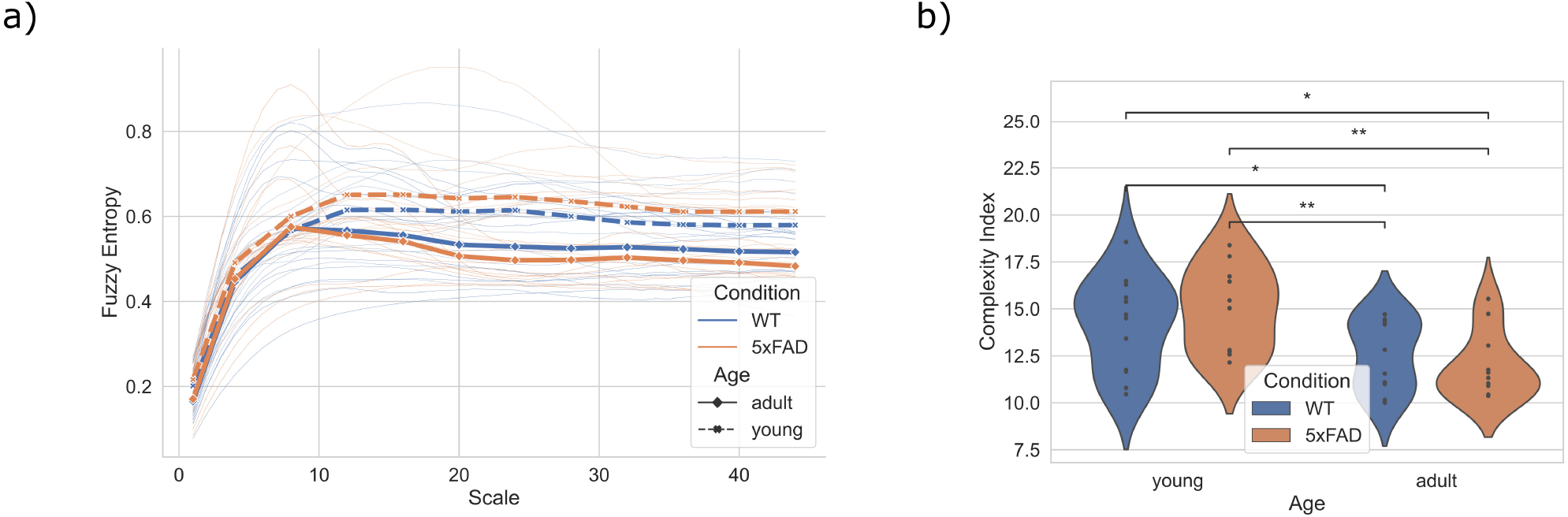
Multiscale entropy (MSE) analysis of *μ*ERG in response to a deterministic stimulus. a) MSE curves of *μ*ERG responses for the CS protocol. Thinner lines: MSE curves of all subjects; thicker lines: average MSE for groups WT-young, WT-adult, 5xFAD-young and 5xFAD-adult. b) Complexity index, calculated as area under the MSE curve. *: *p* < 0.05 and **: *p* < 0.01 for Mann-Whitney U test.

**Figure 5.**
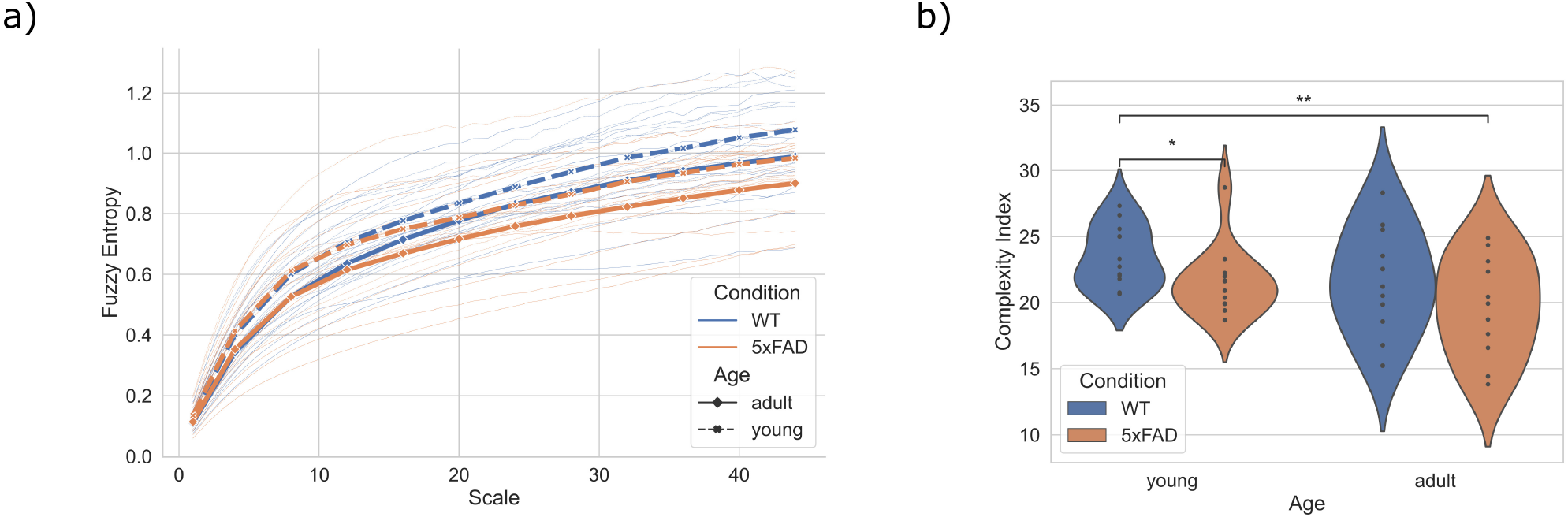
Multiscale entropy (MSE) analysis of *μ*ERG in response to a natural image sequence. a) MSE curves of *μ*ERG responses for the NI protocol. Thinner lines: MSE curves of all subjects; thicker lines: average MSE for groups WT-young, WT-adult, 5xFAD-young and 5xFAD-adult. b) Complexity index, calculated as area under the MSE curve. *: *p* < 0.05 and **: *p* < 0.01 for Mann-Whitney U test.

**Table 1.** Complexity index for the four groups in response to two stimuli

| Stimulus | WT-young | 5xFAD-young | WT-adult | 5xFAD-adult |
| --- | --- | --- | --- | --- |
| Chirp | 14.7 ± 3.85 | 15.04 ± 3.82 | 12.82 ± 3.25 | 11.33 ± 2.43 |
| Natural image | 22.72 ± 3.21 | 21.26 ± 2.14 | 21.22 ± 5.32 | 19.92 ± 5.63 |

\*Median ± 25-75% interquartile range.

## Discussion

We performed multiscale entropy (MSE) analyses of microelectroretinogram (*μ*ERG) signals recorded in retinas of healthy and diseased animals of different ages in response to two types of light stimuli. The *μ*ERG signals exhibited non-decreasing entropy values for a wide range of time scales, which is a signature of the dynamics of complex systems^22, 37^. These results illustrate the complex nature of the electrophysiological activity of the retina exposed to high-information visual fields as those encountered in a natural environment. Further, the responses of young, healthy retinas to natural-like stimuli were more complex than those of diseased retinas regardless of age. For deterministic sinusoidal stimuli, the responses of young retinas were more complex regardless of condition. These findings agree with the theory of complexity-loss with aging and disease^24^, though in our measurements age appeared as a more determinant factor in the reduction of complexity.

Our results suggest that complexity differences due to age are more inherent and less responsive to the nature of the stimulus than those observed under disease. The loss of complexity was first postulated as an age-related phenomenon such that impairment of functional components and/or altered coupling between these components would lead to a progressive decline in adaptability to physiologic stress^38^. Since then, mounting evidence in various physiological outputs showed complexity reductions with age and disease^24, 26, 31, 37, 39^. Our measurements of retinal *μ*ERG bring additional evidence of age-related loss of complexity, a feature that, in this case, emerged for diverse visual inputs. We also observed reductions of complexity in diseased retinas, however, detecting it via MSE analysis required a more complex stimulus that triggered a more complex response in all retinas (21.65 and 13.23 median complexity for NI and CS protocol, respectively). Thus, for entropy tools to be used in early AD diagnosis, it is first necessary to find visual stimuli that elicit complex responses, as it may be the case for white noise inputs^40^.

The MSE curves calculated for the retinal responses showed entropy values close to zero for very small scales, implying that *μ*ERG signals had no significant variability at these scales. Variations at small time scales correspond to high frequency components, and since the signals were low-pass filtered at 30 Hz, low entropy values for time scales < ⌈200/30⌉ = 7 were expected. Further, retinal *μ*ERG seems to function at temporal scales <~ 10 Hz^11^, supporting the notion of low variability at lower scales. In addition, for *FuzzyEn* calculation we used sequences of length 2 (*m* = 2), in agreement with previous work^41^, due to the limited length of the signals. We also used a distance threshold of 0.2 times the standard deviation of signals (*r* = 0.2), a common value used in previous reports^41^. However, we note that *FuzzyEn* changes smoothly with *r*^36^, and in fact, our MSE curves did not substantially change with *r* between 0.1 and 0.5.

In our experiments, small pieces of the retina were placed on microelectrodes to capture, after low-pass filtering, the overall contribution of photoreceptors, bipolar cells, and amacrine cells. Previous work showed that ERG responses in 5xFAD retinas were altered at certain ages^20^. More specifically, 5xFAD retinas older than 6 months-old exhibited reductions, as compared to healthy ones, in traditional ERG features, namely P3, P2 and OP, associated with photoreceptors, bipolar cells and amacrine cells, respectively. However, no changes were detected at 6 months-old (the youngest age analyzed in this study), although a retinal ganglion cell-mediated response was reduced at this age. This indicates that amplitude-based tools are able to detect physiological changes in the retina only during advanced stages of the disease. Conversely, in our measurements we observed significant reductions in the complexity of 5xFAD retinas as compared to healthy ones for the younger group (*i.e*., 2-3 months-old), whereas for the older group (i.e. 4-6 months-old) the differences were not significant. This suggests that MSE analyses may be able to detected physiological changes early in the development of the disease that go otherwise undetected by traditional amplitude-based tools.

Our findings support the theory of complexity-loss with aging and disease and demonstrate that MSE analyses may be a valuable tool to detect physiological changes in senescence or during illness. Our results also present additional evidence of the electrophysiological changes in the retina of an transgenic animal model of AD and can have great implications for early AD diagnosis. Future work should determine whether similar effects are observed in AD patients, and what type of visual stimuli should be used to elicit retinal behaviors with enough complexity to detect differences.

## Methods

The data analyzed in this work were described with more details in a previous publication of our group^21^, and thus we provide here only a brief description of how the recordings were obtained. The entropy/complexity analyses of *μ*ERG signals presented here have not been the subject of previous analyses or publications, and thus we provide a more detailed description of the entropy tools used in this work.

### Alzheimer’s animal model

5xFAD mice (Jackson laboratory, Bar Harbor, Maine, USA) and wild-type (WT) mice (B6SJLF1/J) were used in this study^21^. These animals were maintained in Universidad de Valparaíso, with a time routine of 12:12 hrs light/dark cycle, with controlled temperature, and water and food *ad libitum*.

The animals were grouped into young subjects (2-3 months-old), age at which the animals start exhibiting amyloid-*β* brain aggregation, and adult subjects (6-7 months-old), which present thorough accumulation of plaques in the brain and behavior alterations42. All experimental procedures followed protocols approved by the bioethics committee of Universidad de Santiago de Chile, following the international guidelines on animal handling and manipulation and the Chilean National Agency for Research and Development (ANID) bioethics and biosecurity standards.

### *μ*ERG recordings from mice retina

The protocol for *μ*ERG recordings was performed using a Multi-Electrode Array (MEA) (USB256, Multichannel Systems GmbH, Reutlingen, Germany) with 252 electrode channels and a sampling rate of 20 kHz, which recorded the responses of a small piece of the retina (*n* =13, 11, 12 and 11 for WT-young, WT-adult, 5xFAD-young and 5xFAD-adult, respectively). In brief, all the recordings were stored in a computer for offline analysis. Before the experiments, the animals were dark-adapted for 30 min, and then profoundly anesthetized with Isofluorane (Baxter, Deerfield, Illinois, USA) and euthanized. Eyes were removed and enucleated under dim red light, and eyecups were maintained in Ames medium (Sigma-Aldrich, San Luis, Missouri, USA) at 32 °C and pH 7.4 and oxygenated with a mixture of 95% O2 & 5% CO2. Small pieces of the retina were positioned on an o-ring (MWCO-25000, Spectrumlabs, Rancho Dominguez, California, USA) with polylysine treatment (Product P4707, Sigma-Aldrich, San Luis, Missouri, USA).The retina was stimulated with different visual stimuli created on MATLAB software (Natick, Massachusetts, USA), projected on it using a LED projector (PB60G-JE, LG, Soul, South Korea), mounted in an inverted microscope (Eclipse T200, Nikon, Minato, Tokyo, Japan). The average radiance of each stimulus was 70 nW/mm^2^ (Newport Corporation, Irvine, California, USA) with a spectral content ranging between 460 nm and 520 nm (Ocean Optics Inc, Dunedin, Florida, USA). All stimuli thoroughly covered the piece of the retina.

We applied two visual stimulation protocols consisting of 21 repetitions each of: 1) a chirp stimulus (CS) protocol, and 2) a natural image (NI) protocol. The CS protocol consisted of an ON-light flash (3s) and an OFF-dark period (3s) followed by a sinusoidal light stimulus of increasing frequency (1 to 10 Hz) and fixed maximum intensity, and followed by a sinusoidal light stimulus of 1 Hz with increasing intensity. Each repetition of the CS protocol lasted ~ 35 s (~ 12 min total). The NI protocol consisted of a sequence of images recorded in natural habitat and presented at a refresh rate of 60 fps^21^. Each repetition of the NI protocol lasted ~ 11 s (~ 5 min total). In the experiments, we used a fotodiode (PDA100A-EC, Thorlabs, Newton, New Jersey, United States) to control the retinal illumination stimulation (Fig. 1).

Selected channels according to a quality criterion21 were averaged to obtain a single *μ*ERG signal for each piece of retina stimulated with the CS and NI protocols. The median number of selected channels (± 25-75% interquartile range) for the WT-young, 5xFAD-young, WT-adult and 5xFAD-adult groups was: 197.5 (±64), 206 (±116), 186 (±78) and 203.5 (±99), respectively. The averaged *μ*ERG signal was LP-filtered at 30 Hz, then subsampled at 200 Hz, and for each stimulation protocol, the recorded *μ*ERG signal was averaged over 21 repetitions.

### Spectrum and coherence

We calculated the power spectral density (PSD) of signals for frequency-domain characterization by means of Welch’s method^43^. This method estimates PSD as an average of the periodogram of multiple overlapping segments, thereby reducing noise in PSD estimation at the cost of frequency resolution. We used a Hann window of size 256 and an overlap of 128. In addition, for the CS protocol, we calculated the coherence, *C_xy_*, between the chirp stimulus and the *μ*ERG response by means of the cross-spectral density, *P_xy_*, and the PSD of each signal:

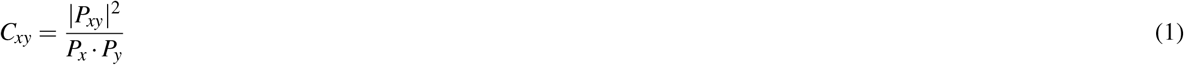

Here, *P_xy_* can be viewed as the PSD of the cross-correlation signal, and it is estimated using Welch’s method. For both PSD and coherence calculation we used the Scipy Python library v1.6.2.

### Multiscale entropy

In this study, we used the concept of entropy as a measure of signal uncertainty or disorder. The sample entropy^44^, *SampEn*, is a regularity statistics so that higher *SampEn* values are assigned to less predictable, more irregular time series. Given a time series *X* of length *N*, *SampEn* counts, for each subset sequence *i* of length *m*, the number of sequences of the same length that are at a distance smaller than *r* from the sequence *i*. Let this number be 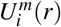, then:

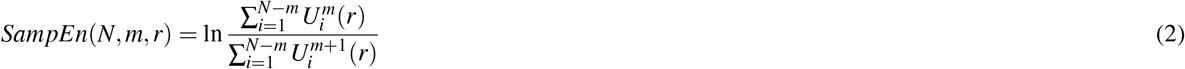

Here, 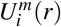 excludes the comparison of sequence *i* with itself. *SampEn* overcomes some difficulties of other entropy measures, namely, the need of many samples, and sensitivity to noise. *SampEn* resembles the so-called approximate entropy, *ApEn*, but it is a less biased statistic since, differently to *ApEn*, it eliminates the self-matching in the counting process. However, *SampEn* shows high sensitivity to parameter selection since, when comparing a pair of subsets, a Heaviside function is applied as a similarity metric after taking the distance – the contribution of such a pair is 1 if the distance between the subsets is less than *r*, and 0 otherwise. In contrast, the fuzzy entropy (*FuzzyEn*)^36^ uses a fuzzy degree of similarity, 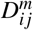, which instead of being binary, depends on the distance between the pairs, 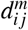, through a continuous, concave function, typically quadratic exponential:

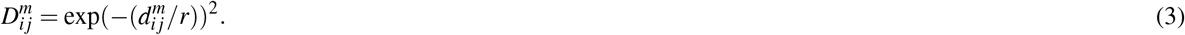

The multiscale entropy (MSE) method was proposed to account for the multiple time scales present in physiological processes^22, 37^. Under this framework, a metric of entropy, *e.g*., the *FuzzyEn*, is calculated for the original time series, i.e., time scale 1, and for coarse-grained time series constructed by averaging as many samples as the current scale, in non-overlapping windows. For example, for time scale 2, the corresponding coarse-grained series is of length *N*/2, and each sample corresponds to the average of two consecutive samples of the original time series. This coarse-graining procedure is illustrated in Fig. 6 for two types of noise and for an electrophysiological signal. The coarse-grained time series appears less irregular for larger scales due to averaging for uniform noise. However, pink noise (a complex signal example) and the example *μ*ERG signal exhibit a degree of irregularity that appears to persist across all scales.

**Figure 6.**
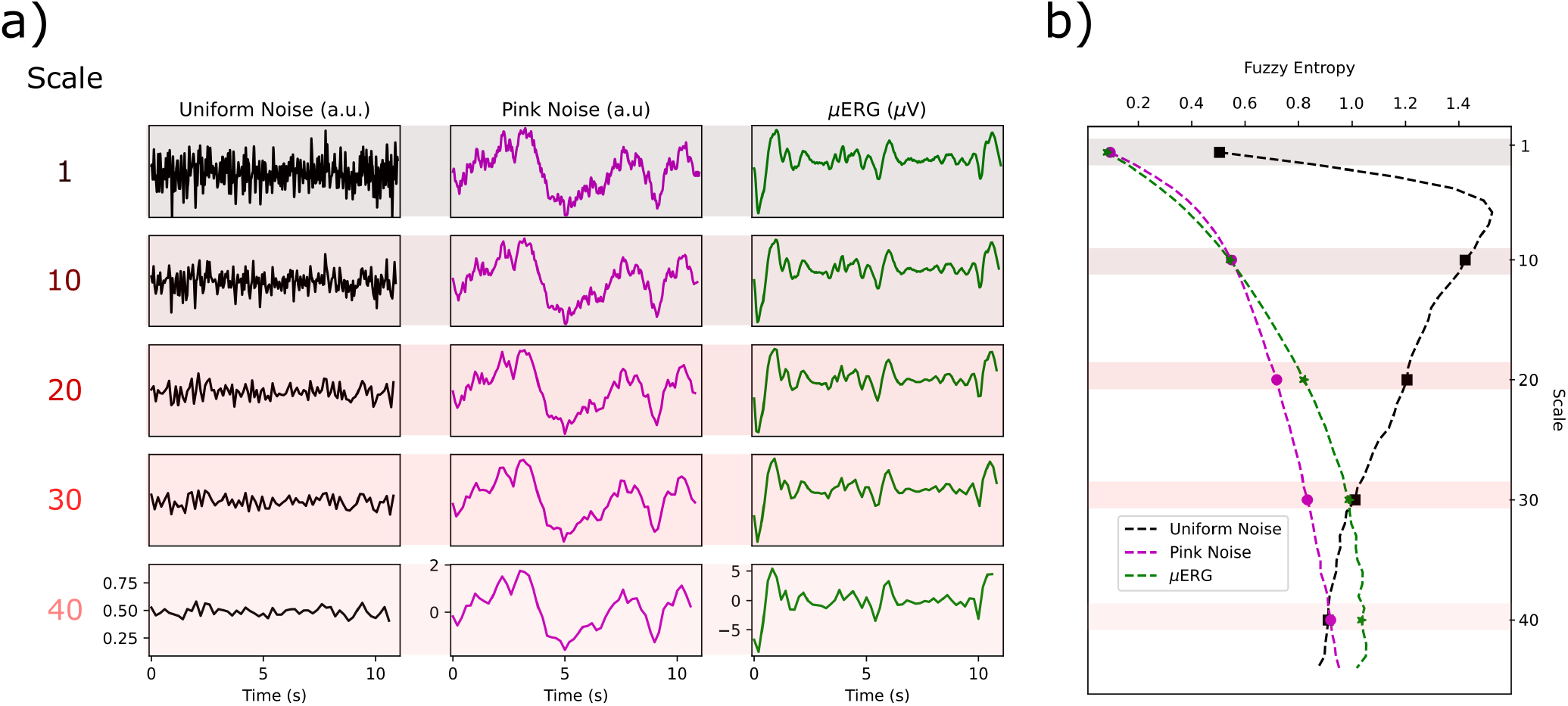
Illustration of the multiscale entropy (MSE) procedure. a) Effect of the coarse-graining procedure on uniform noise (left), pink noise (middle) and an example *μ*ERG signal (right). Original signals (scale 1, top) were low-pass filtered at 30 Hz (sampling frequency 200 Hz), and the coarse-grained signals were calculated for four different scales: 10, 20, 30 and 40. The irregularity of the uniform noise signal decays for scale > 10, whereas for the other two signals, the main irregular features persist across all scales, reflecting a more complex underlying structure. b) MSE curves calculated for the signals in a). The entropy for the scales shown in a) are shaded. Note that for scales > 10, the entropy of uniform noise decays monotonically, whereas for pink noise and the *μ*ERG signal, it increases.

Note that when coarse-graining a time series for a given scale *τ*, the resulting reduced series depends on the choice of the starting position for the initial window, so that there are *τ* possible coarse-grained resulting series. In the composite MSE (CMSE), the entropies of all these coarse-grained time series corresponding to scale *τ* are calculated and then averaged to obtain a single entropy value for *τ*. This procedure ensures more reliable results for larger scales^45^. Further, in the refined CMSE (RCMSE), the logarithm for entropy calculation is taken after computing the number of similar sequences, which reduces the probability of obtaining undefined entropy and increases the accuracy of entropy estimation^46^. In this study, we calculated the RCMSE of *μ*ERG responses with the *FuzzyEn* using the Neurokit2 toolbox for Python^47^. For the NI protocol, the shortest one, each *μ*ERG response comprised 2,200 samples. Since the *FuzzyEn* demonstrated strong consistency in entropy estimation with a time series of 50 samples or greater^36^, we calculated the RCMSE up to scale 2200/50 = 44. For *FuzzyEn* calculation, we used *m* = 2 and *r* = 0.2.

### Complexity

More complex signals have consistently higher entropy values across different time scales^22, 37, 48^, and accordingly, the area under the MSE curve has been used to quantify complexity^27, 31, 37^. We calculated a complexity index, *C_i_*, as the cumulative sum of the RCMSE for scales 20 – 44, noting that 1) at the time scale of 20 and sampling frequency of 200 Hz, frequency components should concentrate to ≤ 10 Hz, 2) *μ*ERG responses strongly attenuate for sinusoidal light stimuli > 10 Hz^11^, and 3) coherence for the CS protocol decayed for frequencies of ~ 10 Hz and above.

### Statistical analysis

For the complexity index calculated for the four groups, WT-young (*n* = 13), WT-adult (*n* = 11), 5xFAD-young (*n* = 12) and 5xFAD-adult (*n* = 11), we performed group-to-group comparisons by means of the Mann-Whitney U rank test. Significance level was set at *α* = 0.05.

## Data availability

The data analyzed during the current study are available in the Zenodo repository, DOI: 10.5281/zenodo.5798072.

## Acknowledgements

This work was partially supported by grants ANID/FONDECYT 11190822 (L.E.M.), 1210622 (C.D.-A.) and 1200880 (A.G.P.), ANID/Millennium Science Initiative Program NCN19_161 (L.E.M.), ICM-P09-022-F (A.G.P.), and grants 2018-AARG-591107, ANID/FONDEF ID20I10152, ANID/Anillo ACT210096 (C.D.-A.).

## Author contributions statement

J.A.-A, A.G.P., M.C. and L.E.M. designed the study; J.A.-A performed the experiments; C.D.-A. provided all animals; L.E.M, C.R., S.G., and M.C. analyzed the data; L.E.M., J.A.-A and C.R. wrote the first draft of the manuscript. All authors revised and approved the manuscript.

